# Highly accurate barcode and UMI error correction using dual nucleotide dimer blocks allows direct single-cell nanopore transcriptome sequencing

**DOI:** 10.1101/2021.01.18.427145

**Authors:** Martin Philpott, Jonathan Watson, Anjan Thakurta, Tom Brown, Tom Brown, Udo Oppermann, Adam P Cribbs

## Abstract

Droplet-based single-cell sequencing techniques have provided unprecedented insight into cellular heterogeneities within tissues. However, these approaches only allow for the measurement of the distal parts of a transcript following short-read sequencing. Therefore, splicing and sequence diversity information is lost for the majority of the transcript. The application of long-read Nanopore sequencing to droplet-based methods is challenging because of the low base-calling accuracy currently associated with Nanopore sequencing. Although several approaches that use additional short-read sequencing to error-correct the barcode and UMI sequences have been developed, these techniques are limited by the requirement to sequence a library using both short- and long-read sequencing. Here we introduce a novel approach termed single-cell Barcode UMI Correction sequencing (scBUC-seq) to efficiently error-correct barcode and UMI oligonucleotide sequences synthesized by using blocks of dimeric nucleotides. The method can be applied to correct either short-read or long-read sequencing, thereby allowing users to recover more reads per cell and permits direct single-cell Nanopore sequencing for the first time. We illustrate our method by using species-mixing experiments to evaluate barcode assignment accuracy and evaluate differential isoform usage and fusion transcripts using myeloma and sarcoma cell line models.

## Introduction

Single-cell RNA sequencing (scRNA-seq) is a widely adopted method for profiling the transcriptome in biology and medicine [1]. Current scRNA-seq methods can be broadly classified as well-based or droplet-based. Well-based methods, such as SMART-seq2/3 [2, 3], partition single cells into individual wells of a multi-well plate which act as a discrete reaction vessel for subsequent library production, followed by short-read sequencing. SMART-seq2/3 has the advantage of producing full-length transcripts, although inferring individual transcripts from short-read sequencing remains challenging. Furthermore, this method is limited to processing hundreds of cells at a very high cost per cell. Droplet-based methods, such as Drop-Seq, InDrops or commercial solutions such as 10X Genomics Chromium [4-6], co-capture cells and oligonucleotide-barcoded RNA-capture microbeads in droplets within an oil emulsion. Each droplet becomes a discrete reaction vessel, associating a different barcode with each cell’s RNA, followed by pooled library production and short-read sequencing. Barcoded RNA-capture microbeads for Drop-seq are manufactured using a manual split and pool process that creates a unique bead barcode region within the oligonucleotide capture sequence [4]. These droplet-based methods are capable of reporting on many thousands of cells at a dramatically reduced cost per cell but are only capable of reporting on the 5’ or 3’ ends of transcripts when using Illumina sequencing.

Long-read sequencing platforms, such as Oxford Nanopore [7], have the potential to deliver the throughput and economy of droplet-based short-read sequencing methods. In addition, it provides true end-to-end sequencing of transcripts, therefore allowing examination of RNA splicing events, single nucleotide polymorphisms, structural variation, imprinting and measurement of chimeric transcripts at the single-cell level. However, the drawback of Nanopore sequencing is its high error rate compared to Illumina-based short-read sequencing (5-15% vs <1%) [8]. For many bulk applications, the advantages of long reads outweigh the low-read accuracy, which can be overcome with consensus sequences from homogenous samples. However, in scRNA-seq, maintaining the fidelity of the barcode and UMI region is indispensable, and accordingly has hampered the adoption of single-cell Nanopore sequencing.

We here describe a barcode and UMI error correction method (scBUC-seq), whereby the barcode and UMI regions of the oligonucleotide-barcoded RNA-capture microbeads are synthesized using dual nucleoside phosphoramidite building blocks. Whilst increasing the length of barcode regions by additional rounds of split-and-pool synthesis using single phosphoramidites could possibly provide means for barcode error correction [9], this approach would substantially increase the time and cost of manufacture of these beads, while also reducing the yield. Furthermore, it would be impossible to correct the error rate within the UMI, which is synthesised randomly. Instead, we take the novel approach of building the barcode and UMI region using homodimeric reverse phosphoramidites during the split and pool process. These repeated bases allow error detection and correction of long-read single cell sequencing, without the need for parallel short-read sequencing.

## RESULTS

### Barcode and UMI error correction strategy

In order to ensure highly accurate barcode and UMI assignment, we incorporated homodimeric nucleotides into the synthesis reaction (scBUC-seq; single cell Barcode Umi Correction sequencing) (Figure 1a). This allows for accurate assignment of barcodes to cells that have not been affected by either PCR or sequencing errors (Figure 1b). Additionally, unlike previous computational methods for Nanopore correction, this approach also allows error correction of the UMI to a high degree of accuracy (Figure 1c). In order to correctly assign barcodes to cells, we developed a computational strategy in which true barcodes were identified in a two-pass assignment method. Firstly, we identified true barcodes based on nucleotide pair complementarity across the full length of the barcode. Next, we used these true barcodes as a guide to error correct the remaining barcodes. Using simulated data, we show that our strategy is capable of correcting barcodes with a high sequencing error rate, with 96% of barcodes recovered with a sequencing error rate of up to 10% (Figure 1d). Next, we modified the directional network-based UMI correction method, first proposed by UMI-tools [10], to deduplicate UMI sequences. In simulated data we show that UMIs are ineffectually deduplicated with sequencing error rates > 5% when the oligonucleotide is synthesised using single nucleotides. However, using the dual nucleotide blocks and incorporating the sequencing error rate of the UMI into the deduplication strategy, we demonstrate that, even at a sequencing error rate > 10%, we are able to effectively deduplicate UMI sequences (Figure 1e and Figure 1f, Supplementary Figure 1).

**Figure 1:**
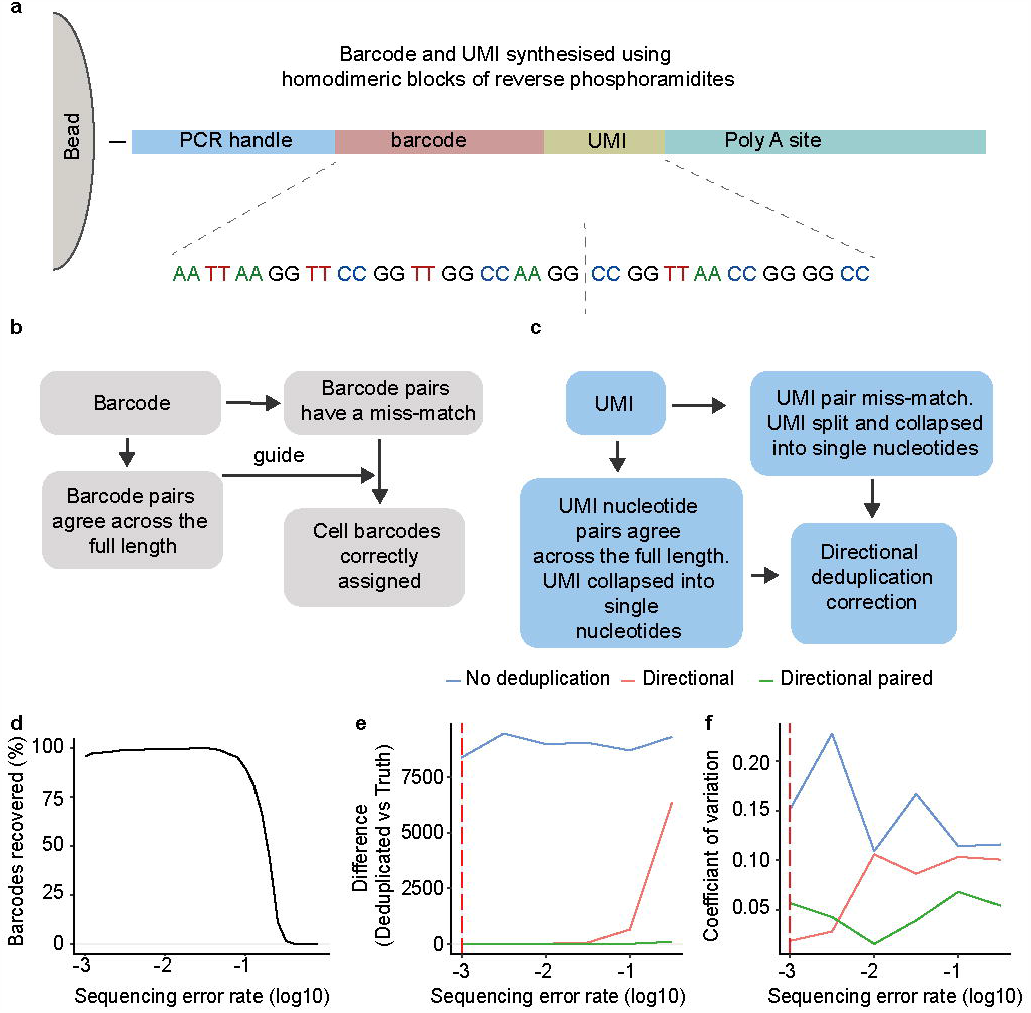
Developing a strategy to error correct barcode and UMI sequences from droplet-based sequencing. **a** Schematic bead and oligonucleotide structure using dimer blocks of nucleotides for BUC-seq. **b** Cell barcode assignment strategy. **c** UMI deduplication strategy. **d** Simulated data showing the number of barcodes recovered with increasing simulated sequencing error rates. **e, f** Simulated data showing the difference and coefficient of variation between the deduplicated UMIs and the ground truth. Deduplication was performed using a basic directional network-based approach and accounting for sequencing errors within paired nucleotides.

### Accurate assignment of cell barcodes and unique molecular identifiers within sequencing data

In order to validate our method, we prepared human HEK293T and mouse 3T3 single-cell Dropseq libraries from approximately 500 cells using the DolomiteBio Nadia microfluidic encapsulation system [11], followed by Illumina short-read sequencing. The low sequencing error rate associated with this technology provided a test bed in which to evaluate the performance of our barcode and UMI correction methodology. Overall, 68% of all barcodes show complete dual nucleotide block complementarity across the full barcode sequence. This suggests that the basecalling accuracy is 98.4%, which, given that this includes errors introduced during library preparation, aligns with the reported accuracy of Illumina sequencing. However, in barcodes that contain at least one sequencing error, the likelihood of seeing more than one sequencing error is increased (Supplementary Figure 2). This reduces the overall basecalling barcode accuracy to 92%. Using those perfectly aligned reads, we next evaluated their ability to correct inaccurately sequenced barcodes. We evaluated the accuracy of our method by measuring the proportion of human, mouse and mixed species cells, identified following increasing the edit distance (i.e. Levenshtein distance [12]) between the error-sequenced and the accurately sequenced barcodes (Supplementary Figure 3). Analysis of uncorrected error-prone barcodes revealed a low number of correctly assigned cells, with the majority of those cells containing a mixed fraction of mouse and human reads (Figure 2a). Using an edit distance of 4 we found that this resulted in accurate assignment of both mouse and human reads (Figure 2b). Overall, using this edit distance we were able to recover an extra 8 % of total reads. Whilst further reads could be recovered using an edit distance of 6, this was obtained at the expense of increased mixed cells (Supplementary Figure3i).

**Figure 2:**
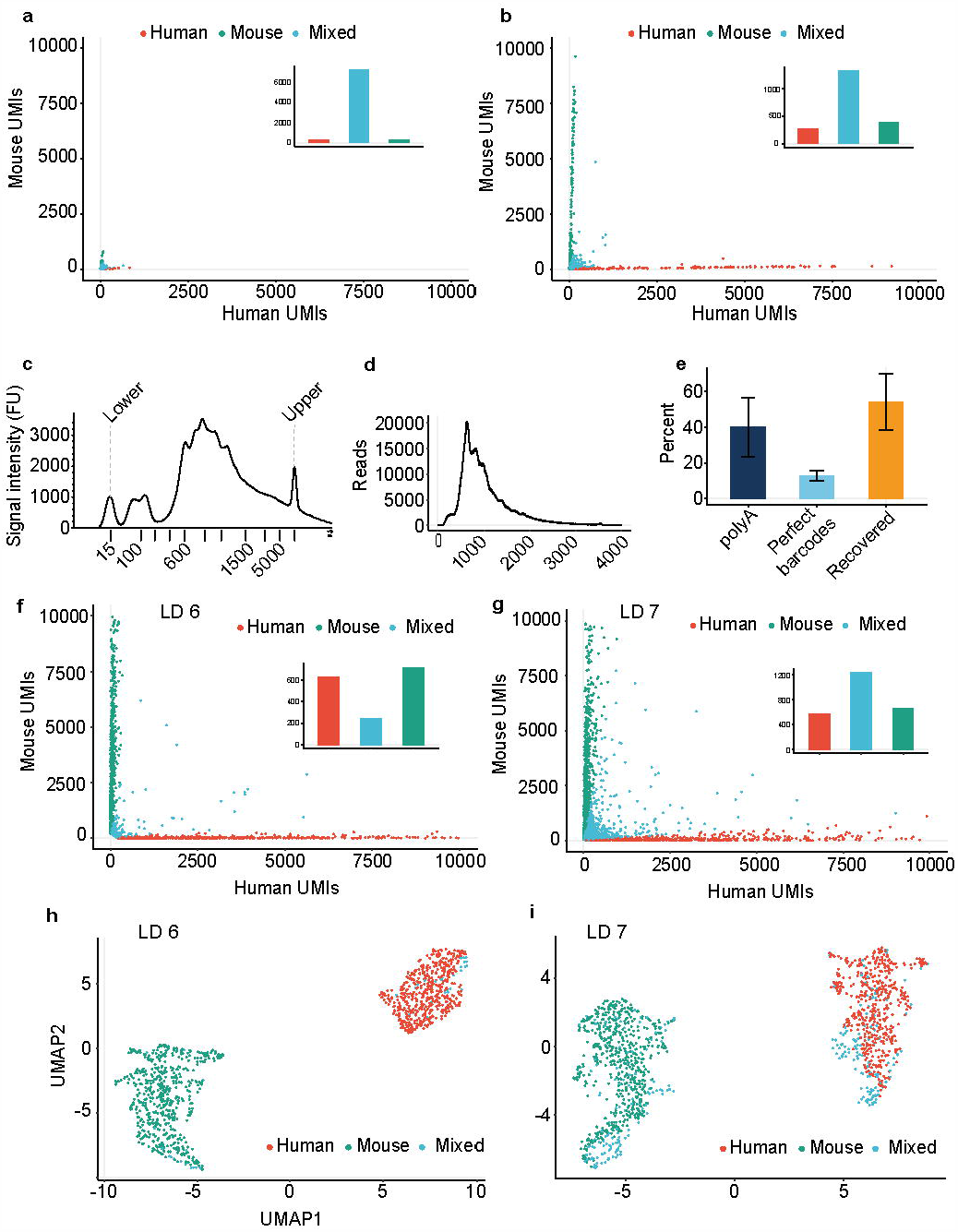
Error correction of both Illumina and Nanopore droplet based scRNA-seq data. Human HEK293T and mouse 3T3 were mixed at a 1:1 ratio and approximately 500 cells were taken for encapsulation and cDNA synthesis. Barcodes and UMIs identified as having at least one sequencing error were processed **a** before and **b** after barcode error correction. The proportion of mouse and human UMIs are shown in the Barnyard plot. Insert bar plots show the number of cells identified for each species. **c** The length of the input cDNA Nanopore library, as measured using a tapestation. **d** The read length of the sequenced Nanopore library. **e** The percent of reads that have a polyA tail. The percent of polyA^+^ reads that show perfect based on the nucleotide pairing complementarity and the percent of reads that ccan be recovered using an Levenshtein distance of 6. Boxes and error bars indicate the means and standard deviations for n=4 individual experiments. Barnyard plots showing the expression of mouse and human UMIs using a **g** Levenshtein distance (LD) of 6 and a **h** Levenshtein distance of 7. The insert bar plots show the number of cells recovered for each species. UMAP plots of the showing human, mouse or mixed human and mouse cells when barcodes are corrected using a i Levenshtein distance of 6 or a j Levenshtein distance of 7.

**Figure 3:**
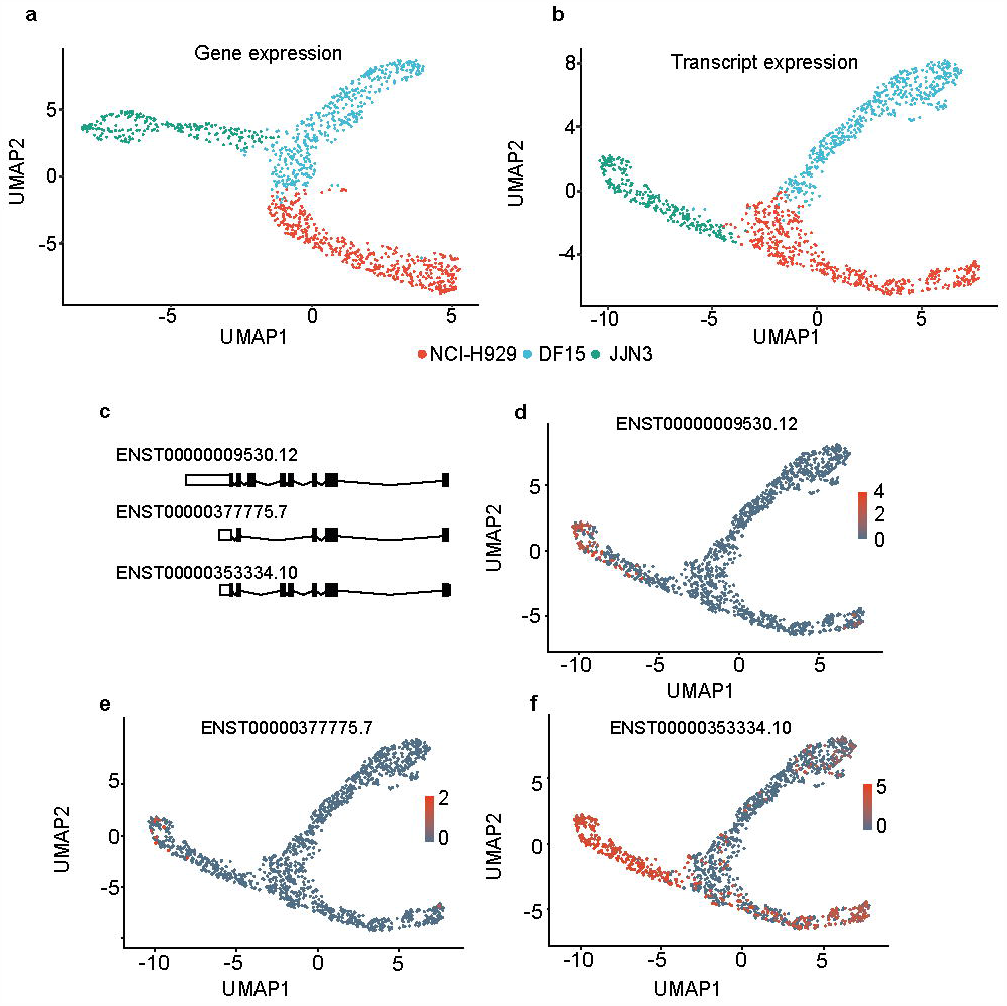
Nanopore droplet based scRNA-seq identifies isoform diversity. NCI-H929, DF15 and JJN3 myeloma cell lines were mixed at a 1:1:1 ratio and approximately 1200 cells were taken for cDNA synthesis and sequenced using a PromethION flow cell. UMAP plot of **a** gene expression and **b** transcript isoform expression. **c** Principal CD74 (HLA-DR) splice variants showing all protein coding transcripts. UMAP plot showing the isoform expression of detected CD74 (HLA-DR) transcripts **d** ENST00000009530.12, **e** ENST00000377775.7 and **f** ENST00000353334.10.

Having demonstrated that our single-cell oligonucleotide design could provide reliable base-calling and barcode error rate information using Illumina data, we next applied the technology to the Oxford Nanopore sequencing platform [7]. Using the same cDNA (Figure 2a and Figure 2b), we found that Nanopore sequencing produced a read distribution of a similar length to that of the input cDNA (Figure 2c ; Figure 2d). We identified the presence of a polyA sequence in 40% (range 24% – 62% over 4 independent experiments) of all Nanopore sequencing reads and detected 12.9% (range 9% − 15%) of these reads showing dual nucleotide complementarity across the full barcode sequence (Figure 2e). This suggests the basecalling accuracy of single-cell nanopore sequencing is 91.8%. Similar to Illumina sequencing, we also observe that if a barcode contains at least one sequencing error there is an increased likelihood of it containing more than one error (Supplementary Figure 2b). Therefore, the overall basecalling barcode accuracy is measured at 86%. We next evaluated the ability of our method to error correct barcodes containing sequencing errors. We found that using a barcode correction edit distance of 6 led to the recovery of 54% (range 43% − 68%) of barcodes containing sequencing errors (Figure 2f, Supplementary Figure 4). Increasing the edit distance to 7 increased recovery to 82% (range 79.8% − 83.6%), however this increased recovery was at the expense of increased numbers of mixed cells (Figure 2g and Supplementary Figure 4c). Despite the increased presence of mixed cells, filtering removed a substantial proportion of these cells and we were still able to observe clear mouse and human cell population separation. Despite this, we opted for the conservative edit distance of 6 for all subsequent analyses.

**Figure 4:**
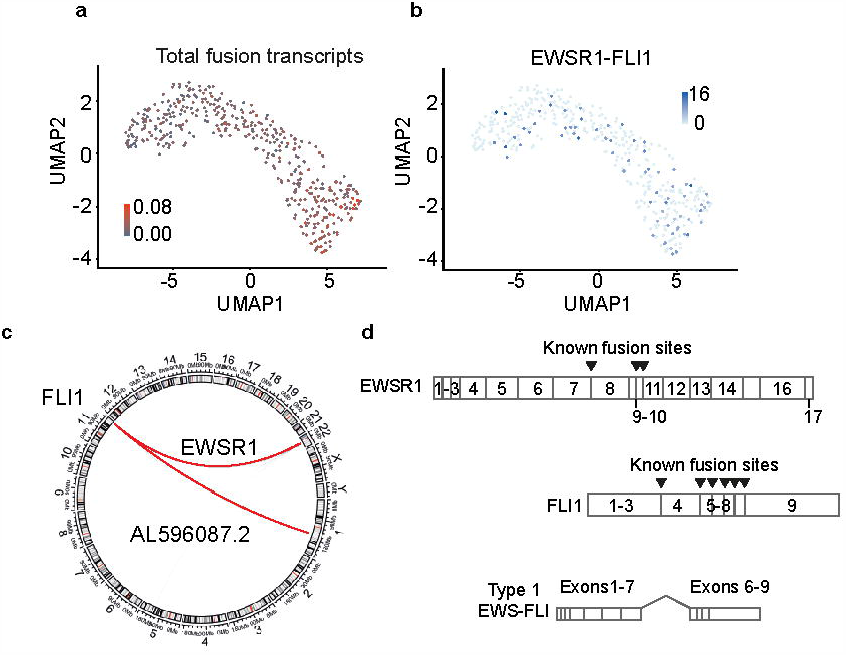
Nanopore scRNA-seq reveals fusion transcripts in Ewing’s cells. A UMAP plot of **a** total fusion transcripts in Ewing’s cells mapped as a parentage of the cell’s total RNA and a UMAP plot showing the expression of the **b** EWS-FLI fusion transcript. **c** A circular representation of the fusion transcripts identified between FLI1 and EWSR1. **d** A schematic showing the structure of the EWSR1 and FLI1 genes. The EWS-FLI1 fusion transcript consists of the 5’ end of the EWS gene and the 3’ end of the FLI1 gene. Arrowheads denote known fusion events and the most common type 1 fusion transcript is shown.

### Identification of alternative transcript isoform usage using scBUC-seq in multiple myeloma cell lines

We applied scBUC long-read sequencing to a mixture (1:1:1 ratio) of NCI-H929, JJN3 and DF15 myeloma cell lines and performed cDNA synthesis on approximately 500 cells sequenced using a MinION device and 1200 cells sequenced using the PromethION platform. Following filtering (Supplementary Figure 5 and Supplementary Figure 6), we show that Nanopore sequencing is capable of resolving the different myeloma cell types at both the gene level (Figure 3a and Supplementary Figure 7a) and the transcript level (Figure 3b and Supplementary Figure 7b). There was also a good correlation between Nanopore and Illumina gene counts (Supplementary Figure 8). We next searched for differentially expressed transcripts between cell types and clusters. In this experiment we observed cell-type specific usage for 359 genes and 416 differentially expressed isoforms. Differential transcript usage was particularly apparent in the marker CD74 (Figure 3c; Figure 3d; Figure 3e; Figure 3f), which is a potential therapeutic target in multiple myeloma [13]. Furthermore, in agreement with the literature and the biology of plasma cells [14], we observed a significant differential expression of both immunoglobulin kappa and lambda light-chain isoform usage between the different Myeloma cell lines (Supplementary Figure 9 and Supplementary Figure 10).

### Identification of fusion transcripts using long-read single-cell sequencing

We next examined how scBUC long-read sequencing performs at measuring fusion transcripts as a result of genomic translocations. Long-read sequencing permits the analysis of chimeric sequences between two different gene regions that are drivers of certain cancers. To illustrate the principle, we selected Ewing’s sarcoma which harbours such oncogenic fusion protein drivers. The most common chromosomal translocation in Ewing’s sarcoma is the t(11:22)(q24:q212) translocation, which generates the EWS-FLI protein, consisting of a fusion between EWSR1 (Ewing’s sarcoma breakpoint region 1) and the Ets transcription factor FLI1 (Friend leukemia integration 1) gene [15]. As a consequence of this fusion, the EWS-FLI protein regulates the expression of numerous target genes that maintain stem cell phenotypes, promote cell proliferation, survival and drug resistance [16, 17]. We performed scBUC-seq on STA-ET-1 Ewing’s cells, which are known to express the EWS-FLI protein, and measured the presence of this fusion transcript within each single-cell. Given that fusion transcripts can be falsely detected as a consequence of PCR artefacts [18], we first used the mixed-species data (Figure 2f) to determine the frequency of false-positive fusion events. We show that 5% of total reads contain a fusion event, with 35 % of these reads showing mixed human and mouse fusion transcripts (Supplementary figure 11). This suggests that over 70% of detected fusion transcripts could be registered as a result of PCR artifacts. However, application of a filtering threshold based on a minimum of 5 UMIs for each fusion event removed all of the mixed human/mouse fusion reads (Supplementary figure 11b and Supplementary figure 11c). This strategy suggests that a filter of 5 UMIs is sufficient to eliminate the vast majority of false positive fusion events and was therefore used to filter our Ewing’s cell data. Following filtering, we detected a total of 10258 unique fusion transcripts allowing measurement of the EWS-FLI fusion protein in 17% of cells (Figure 4a and Figure 4b). In addition, we observed a novel fusion transcript between FLI1 and the long non-coding RNA AL596087.2 (Figure 4c). Several EWS-FLI fusion transcripts have been reported [19]. We accurately detected the presence of the most common “type 1” form in our single-cell data, consisting of the first seven exons of EWSR1 joined to exons 6-9 of the FLI1 gene (Figure 4d and Supplementary Figure 12).

## DISCUSSION

Recent advancements in single-cell droplet-based sequencing technologies have enabled molecular profiling of the transcriptional status of cells and tissues at the single-cell level. However, transcriptional activity is typically summarised at the gene-level due to the limitations of short-read sequencing technologies that typically only allow sequencing of the 3’ portion of a gene transcript. The recent development of long-read sequencing technologies such as PacBio single-molecule real-time (SMRT) sequencing or Oxford Nanopore sequencing promises to revolutionise the sequencing of full-length transcripts [20]. However, its application to single-cell experiments has been stymied by the high base-calling error rates associated with long-read sequencing technologies. This makes it challenging to simultaneously assign a sequencing read to a cell and correct for library-associated PCR duplication errors.

SMRT sequencing has addressed this using circular consensus sequencing (CCS), where subreads are generated by multiple passes around a circularized template during sequencing, allowing a consensus sequence to be determined with >99% accuracy [21]. Although this approach limits the effective read length to <15 kb, which would be an important consideration for some long-read sequencing applications, this would be sufficient to sequence almost all polyA transcripts for single-cell long-read sequencing. However, analysis of single-cell data is highly dependent on acquiring sufficient reads per cell, typically 30,000-100,000 reads per cell. Currently, the highest capacity PacBio SMRT cell returns a maximum of 4 million reads, meaning a single run could report on just 40-133 cells at a prohibitively high cost per cell. Recently, Zeng et al described high-throughput single-cell isoform sequencing (HIT-scISOseq) [22], which, by concatenating multiple full-length cDNAs into a single insert, was able to return 10 million reads from a single SMRT cell. They used HIT-scISOseq to distinguish between a mixture of thousands of injured and uninjured corneal epithelial cells, although were only able to detect an order of magnitude less transcripts per cell than matching short-read scRNAseq. It is unclear how this method would fare with more complex populations of cells, where it would likely be necessary to reduce the number of cells per run into the range of a few hundred cells to reach sufficient transcript coverage. On the other hand, Oxford Nanopore sequencing is more scalable, with a range of flow cells suitable for hundreds to thousands of cells per run at 40,000 reads per cell, as well as the ability to run multiple flow cells in parallel, at a cost per cell that can rival or beat Illumina short-read sequencing, making this approach attractive for single-cell long read sequencing.

Several groups have reported using short-read Illumina sequencing to error correct long-read Nanopore single-cell sequencing [23-25]. While this approach was able to increase assignment rates from just ∼6% to >60%, the requirement to independently construct and sequence two libraries considerably raises the cost of single-cell sequencing. Moreover, accurate UMI assignment is challenging with this approach because of the random nature of the UMI generation and the low base-calling accuracy of Nanopore sequencing. Volden et al used a Rolling Circle Amplification to Concatemeric Consensus (R2C2) method to error correct Nanopore sequencing [26]. Although this method achieved 96% sequencing accuracy, this still only translated to 72% of barcodes demultiplexing correctly, with 45% of UMIs not matching against parallel Illumina sequencing. Furthermore, the increased read length needed to support this error correction approach is prone to increased error rates for longer reads in the late stages of a sequencing run [27].

By modifying the barcode and UMI synthesis methodology, we build our oligonucleotide sequences using homodimeric reverse phosphoramidites during the split and pool process. Having a homodimer provides us with the ability to detect base-calling errors within both the barcode and UMI sequences. We use the highly accurate barcodes, determined by full complementarity of the blocks of homodimer nucleotides across the full oligonucleotide length, as a guide to error correct barcodes with sequencing errors. Furthermore, we adapt directional network approaches, first published by UMI-tools [10], to account for errors within the UMI sequence. Using this approach, we can effectively error correct single-cell sequencing barcodes, with over 80% recovery of our reads when using an edit distance of 7, or over 60% recovery when using a conservative edit distance of 6. Furthermore, we can also deduplicate UMIs with a high level of accuracy. Our approach has multiple advantages over current methodologies to correct error-prone sequencing. First, our approach uses direct Nanopore sequencing, which circumvents the need for additional short-read alignment data. Second, we are able to provide a base-calling accuracy rate for each barcode and UMI sequence and, by using the combined accuracy of all barcodes and UMIs, we can approximate the overall accuracy of a single-cell sequencing experiment. The barcode and UMI sequencing accuracy information is then applied to recover single-cell barcodes and deduplicate UMI sequencing. We show that this approach can be used to error-correct both short-read (Illumina) and long-read Nanopore sequencing data, thereby recovering sequencing data that would otherwise be lost due to barcode mis-assignment.

While we have shown that our method can error-correct sequencing errors to a high accuracy, the method could be further improved by synthesising our oligonucleotide structure in blocks of trimeric phosphoramidites, which would also make the computational analysis less complicated. We are currently exploring this, however the synthesis of reverse trimer phosphoramidites is more complex than dimers [4, 6].

Our direct long-read single-cell sequencing technology has the potential to open new avenues within genomics. For example, we demonstrate that it is possible to measure fusion events in chimeric reads, which is only possible with long-read technology. The same approach could also be applied to study biomedically important events such as immunoglobulin (IgG), T-cell receptor (TCR) and B-cell receptor (BCR) gene rearrangements and cellular repertoires. It should be noted that chimeric reads can be generated during the PCR amplification steps of library preparation [18, 28]. We chose to only consider a fusion transcript if the UMI count was greater than 5. However, even with stringent UMI filtering, artefacts may still be present within the data. Therefore, results should ideally be validated with orthogonal methods such as fluorescence *in situ* hybridization (FISH) based techniques if trying to ascribe biological function.

Single-cell long read technology provides detection of full-length transcripts and expands the toolbox of functional genomic techniques, including epigenetics and mutational analyses. Applications of these techniques, along with single-cell copy number variation and mutational analysis, would have a significant potential in diagnostics and understanding of human disease.

## MATERIAL AND METHODS

### Cell lines and reagents

HEK293T, JJN3, H929 and 3T3 cells were purchased from ATCC. DF15 cells were a kind gift by Celgene (now Bristol Myers Squibb). Cell lines were cultured in DMEM low glucose medium supplemented with FBS for no longer than 20 passages. They were mycoplasma tested routinely and authenticated by STR during the course of this project.

### Oligonucleotide synthesis

Bead functionalization and solid-phase phosphoramidite oligonucleotide synthesis was performed by ATDBio. Toyopearl HW-65s resin, purchased from Tosoh Biosciences (product number: 0019815), was used as the solid support. Prior to oligonucleotide synthesis, the initial loading of hydroxyl groups on the resin was reduced via a capping reaction. Capping was performed by suspending the resin in a mixture of acetic anhydride and lutidine in THF, and N-methyl imadazole in acetonitrile for 24 hours. Following capping, the synthesis was performed using an ABI394 DNA synthesiser. The sequence of the capture oligonucleotide is given below:

Bead − 5’ TT − HEG - PC- HEG- TTTTTTTAAGCAGTGGTATCAACGCAGAGTACJJJJJJJJJJJJNNNNNNNNTTTTTTTTTTTTTTTTTTTTTT TTTTTTTT

where ‘J’ indicates a dimer nucleotide added via split and pool synthesis and ‘N’ indicates a degenerate dimer nucleotide.

The barcode was generated via 12 split and pool synthesis cycles [4]. Prior to the first split and-pool synthesis cycle, beads were removed from the synthesis column, pooled and mixed, and divided into four equal aliquots. The bead aliquots were then transferred to separate synthesis columns and reacted with either 3’-DMT-dG-dG-5’-CE, 3’-DMT-dC-dC-5’-CE, 3’-DMT-dA-dA-5’-CE, or 3’-DMT-dT-dT-5’-CE phosphoramidite. This process was repeated 11 times. Following the final split and pool cycle, the resin was pooled, mixed and divided between four columns prior to the synthesis of the UMI and poly-T tail. An equimolar mixture of the four dimer phosphoramidites was used in the synthesis of the degenerate UMI region.

Following the synthesis, the resin was washed with acetonitrile and dried prior to deprotection in aqueous ammonia (55°C, 6 hours).

Reverse directionality dimer phosphoramidites required for the split and pool and UMI region, were purchased as a custom product from ChemGenes: 3’-DMT-dA(N-Bz)5’-Phosphate-3’-dA(N-Bz)-5’-CE, 3’-DMT-dG(N-iBu)5’-Phosphate-3’-dG(N-iBu)-5’-CE, 3’-DMT-

dC(N-Ac)5’-Phosp hate-3’-dC(N-Ac)-5’-CE, 3’-DMT-dT 5’-Phosphate-3’-dT-5’-CE. Reverse directionality monomer phosphoramidites, used for the SMART primer binding site and poly-T tail, were purchased from LGC Link: 3’-DMT-dA (N-Bz)-5’-CE, 3’-DMT-dG (N-iBu)-CE-5’, 3’-DMT-dC (N-Ac)-5’-CE, 3’-DMT-dT-5’-CE (Item numbers: 2022, 2021, 2023, 2020). The modified phosphoramidite reagents were purchased from LGC Link: Spacer-CE Phosphoramidite (Item number: 2129).

### Simulated barcode data

We simulated barcode sequences with a length of 24 (12 blocks of nucleotides pairs) and then imitated the process of randomly introducing PCR errors and sequencing errors into 95% of our barcodes. We then performed a two-pass barcode assignment strategy in which true barcodes were identified based off the nucleotide pair complementarity across the full length of the barcode. These true barcodes were then used as a guide to correct the remaining barcodes based on approximate string matching (Levenshtein distance). The following values were used as values within our simulations unless otherwise stated: Sequencing depth 400; number of UMIs 10-100; barcode-length 24; PCR error rate 1×10^−5^; sequencing error rate 1×10^−1^ – 1×10^−7^ and number of PCR cycles 25.

### Simulated UMI data

We generated simulated UMI data of length 16 (8 blocks of nucleotide pairs) to confirm the accuracy of our UMI correction method by mimicking UMI PCR amplification and sequencing errors seen with Nanopore sequencing. UMIs were generated following an approach that was initially proposed by UMI-tools [10]. Briefly, each UMI was generated at random, with a uniform probability of amplification (0.8-1.0). We simulated PCR cycles so that each UMI was selected in turn and duplicated according to the probability of amplification. PCR errors were added randomly and then any new UMI sequences were assigned new probabilities of amplification. A defined number of UMIs were randomly sampled to simulate sequencing depth and sequencing errors introduced with a specified probability. Finally, we checked for the presence of mismatched double nucleotides within the UMI and if errors were detected, the UMIs were split into two and then separately collapsed into 8bp nucleotides. Unambiguous UMIs were collapsed into 8bp nucleotides without splitting. The number of true UMIs was then estimated from the final pool of UMIs using UMI correction methods proposed in the original UMI-tools manuscript [10]. The following values were used as values within our simulations. Sequencing depth 10-400; number of UMIs 10-100; UMI-length 6 – 16; PCR error rate 1×10^−3^ – 1×10^−5^; sequencing error rate 1×10^−1^ – 1×10^−7^ and number of PCR cycles 4-12.

### Droplet based scRNA-seq

Single-cell capture and reverse transcription (RT) were performed using the Drop-seq approach, as previously described [4]. Briefly, cells were loaded into the DolomiteBio Nadia system microfluidic cartridge at a concentration of 310 cells per microliter. Oligonucleotide beads were synthesised by ATDBio (Oxford, UK). Beads were loaded into the microfluidic cartridge at a concentration of 620,000 beads per mL. Cell capture and lysis were performed according to the Nadia instrument manufacturer’s instructions (DolomiteBio). The droplet emulsion was then disrupted using 1ml of 1H, 1H, 2H, 2H-Perfluoro-1-octanol (PFO; Sigma) and beads were released into aqueous solution. Following several washes, the beads were then subjected to RT. Prior to PCR amplification, beads were washed and then treated with ExoI exonuclease for 45 mins. PCR was then performed using the SMART PCR primer (AAGCAGTGGTATCAACGCAGAGT) and then cDNA purified using AMPure beads (Beckman Coulter). In order to achieve a high concentration of cDNA the input was subjected to 25 cycles of PCR amplification, rather than the 13 stated in the original Drop-seq protocol. Finally, cDNA was quantified using a TapeStation (Agilent Technologies) using DNA high sensitivity D5000 tape before being split for Illumina or Nanopore library generation.

### Single-cell Illumina library preparation for sequencing

Library preparation for Illumina sequencing was performed as previously described [4]. Briefly, purified cDNA was used as an input for the Nextera XT DNA library preparation kit (Illumina). Library quality and size was determined using a TapeStation (Agilent Technologies) High Sensitivity D1000 tape. High quality samples were then sequenced to a minimum of 50,000 reads per cell on a NextSeq 500 sequencer (Illumina) using a 75-cycle High Output kit using a custom read1 primer (GCCTGTCCGCGGAAGCAGTGGTATCAACGCAGAGTAC).

### Nanopore library preparation for sequencing

Full length cDNA samples were prepared using Oxford Nanopore Technologies SQK-LSK-109 Ligation Sequencing Kit, with the following modifications. Incubation times for end-preparation and A-tailing were lengthened to 15 minutes and all washes were performed with 1.8X AMPure beads to improve recovery of smaller fragments. SFB was used for the final wash of libraries. 50 fmol of library were sequenced on either a MinION FLO-MIN106D R9.4.1 flow cell or PromethION FLO-PRO002 R9.4.1 flow cell (Novogene were used as the sequencing service provider), according to the manufacturer’s protocol. Samples sequenced using the MinION platform were usually sequenced across two or three flow cells so that the final sequencing depth was at least 20 million (∼40,000 reads per cell). Samples sequenced using the PromethION were sequenced using one flow cell so that the final read depth was at least 48 million (∼40,000 reads per cell).

### Illumina-based scRNA-seq analysis workflow

The fastq data was processed using a custom written cgatcore pipeline (https://github.com/Acribbs/TallyNN) [29]. We identified ambiguous and unambiguous reads based on the occurrence of dual nucleotide complementarity within the barcode sequence. The unambiguous barcodes were then used to error correct the ambiguous reads by fuzzy searching using a Levenshtein distance of 4 (unless stated in the figure legend). The barcode and UMI sequence for the corrected read pairs were then collapsed into single nucleotide sequences. The resulting fastq files were used as an input for Kallisto (v0.46.1) bustools (v0.39.3) [30], which was used to generate a counts matrix. This counts matrix was used as an input to the standard Seurat pipeline (v3.1.4) [31].

### Nanopore-based scRNA-seq analysis workflow

We performed base-calling on the raw fast5 data to generate fastq files using Guppy (v4.2.2) (guppy_basecaller –compress-fastq -c dna_r9.4.1_450bps_hac.cfg -x “cuda:1”) in GPU mode from Oxford Nanopore Technologies running on a GTX 1080 Ti graphics card. For each read we identify the barcode and UMI sequence by searching for the polyA region and flanking regions before and after the barcode/UMI. Accurately sequenced barcodes were identified based on their dual nucleotide complementarity. Unambiguous barcodes were then used as a guide to error correct the ambiguous barcodes in a second pass correction analysis approach. We performed fuzzy searching using a Levenshtein distance of 6 (unless otherwise stated in the figure legend) and replaced the original ambiguous barcode with the unambiguous sequence. A whitelist of barcodes was then generated using UMI-tools whitelist (umi_tools whitelist --bc-pattern=CCCCCCCCCCCCCCCCCCCCCCCCNNNNNNNNNNNNNNNN --set-cell-number=1000) [10]. This whitelist was used to assess the quality of cells to read count ratio and used as an input for UMI-tools extract. Next the barcode and UMI sequence of each read was extracted and placed within the read2 header file using UMI-tools extract (umi_tools extract --bc-pattern=CCCCCCCCCCCCCCCCCCCCCCCCNNNNNNNNNNNNNNNN --whitelist=whitelist.txt). Reads were then aligned to the transcriptome using minimap2 [32] *(-ax splice -uf --MD -- sam-hit-only --junc-bed*) using the reference transcriptome for human hg38 and mouse mm10. The resulting sam file was converted to a bam file and then sorted and indexed using samtools [33]. The transcript name was then added as a XT tag within the bam file using pysam. Finally, UMI-tools count (*umi_tools count –per-gene –gene-tag=XT –per-cell – double-barcode*), with modifications that allow it to handle oligonucleotide blocks, was used to count features to cells before being converted to a market matrix format. We modified UMI-tools count to handle the double nucleotide UMIs as defined below. This counts matrix was then used as an input into the standard Seurat pipeline.

### UMI error correction

UMI-tools was forked on Github and the counts functionality was (https://github.com/Acribbs/UMI-tools) modified to handle our double oligonucleotide design. Briefly, if a UMI contained at least one sequencing error the UMI was split into two and then separately collapsed into 8bp nucleotides. UMIs that did not contain a sequencing error were collapsed into 8bp nucleotides without splitting. The directional method implemented within the original UMI-tools was then performed to correct UMI sequencing errors.

### Dimensionality reduction and clustering

Raw transcript expression matrices generated by UMI-tools count (for Nanopore data) or kallisto bustools (for Illumina data) were processed using R/Bioconductor (v4.0.3) and the Seurat package (v3.1.4). Gene matrices were cell-level scaled and log-transformed. The top 2000 highly variable genes were then selected based on variance stabilising transformation which was used for principal component analysis (PCA). Clustering was performed within Seurat using the Louvain algorithm. To visualise the single-cell data, we projected data onto a Uniform Manifold Approximation and Projection (UMAP). Cell type determination was performed using clustifyr v1.0.0 to identify correlated gene expression between single-cells and bulk RNA-seq gene lists from the harmonize database [34, 35].

### Differential gene and isoform expression

Differential expression analysis was performed using nonparametric Wilcoxon test on log_2_(TPM) expression values. Differentially expressed genes and transcripts were selected based on the basis of absolute log_2_ fold change of >1 and the adjusted P value of <. 0.05.

### Identification of fusion transcripts

Nanopore reads were aligned to the hg38 genome with minimap2 (-map-ont --MD --sam-hit-only -junc-bed –secondary=no). The splice junction bed file was generated from the Gencode v36 gtf file using paftools, the minimap2 companion software. The sam file was filtered using samtools to remove all non-primary alignment and supplementary alignments (samtools view -F 3328). Chimeric reads were identified based on the SA SAM tag, which lists all other supplementary alignments. All SAM file processing was performed using pysam v0.15.2. Next the SA tag was inspected and assigned to the genomic feature using a BED file containing records of all known coding genes. The SAM record was updated with Ta, Tb, Tc and Td tags, which defines gene positional information from the BED file. Finally, fusion transcripts were annotated with gene information and the barcode information was used to generate a per cell counts for each translocated read. The counts table was then merged with the original transcript Seurat object. Original UMAP embeddings that were calculated for the transcript only level analysis was used for visualisation.

PCR artifacts must be taken into consideration when investigating novel isoforms or translocations. Most PCR duplications and artefacts can be eliminated when the UMI is accounted for, but some artifacts may remain. Reverse transcription (RT) artifacts are a lot more difficult to identify because RT introduces template switching between homologous sequences leading to increased chimeric cDNA [18, 28]. However, to minimise the false positive translocations in our data we used thermostable RT enzyme, we removed exonic chimeric transcripts and required a minimum of 5 UMIs per translocation event.

## Data availability

Sequencing data has been deposited to GEO under accession number GSE162053.

## Code availability

Source data is provided with this manuscript. All custom pipelines used within the analysis is available on Github (https://github.com/Acribbs/TallyNN). Modifications to the UMI-tools code is also available as a fork on Github (https://github.com/Acribbs/UMI-tools).

## Funding

Research support was obtained from Innovate UK (T.Bs, T.Bj, U.O, M.P, A.P.C), the National Institute for Health Research Oxford Biomedical Research Unit (U.O), Cancer Research UK (CRUK, U.O), the Bone Cancer Research Trust (BCRT) (A.P.C and U.O), the Leducq Epigenetics of Atherosclerosis Network (LEAN) program grant from the Leducq Foundation (U.O), the Chan Zuckerberg Initiative (A.P.C) and the Myeloma Single Cell Consortium (A.T, U.O). A.P.C. is a recipient of a Medical Research Council (MRC) career development fellowship (MR/V010182/1).

## Contributions

A.P.C designed and developed the method, implemented and analysed the data. M.P, U.O and A.P.C conceived the study with contributions from T.B Jr, T.B Sr. M.P, J.W and A.P.C performed experiments. M.P, U.O and A.P.C wrote the manuscript with input from all authors. All authors approved the final manuscript.

## Competing interests

A.T is a full-time employee and shareholder of Bristol Myers Squibb. JW, TBj and TBs are shareholders of ATDBio. MP, UO, A.P.C are inventors on patents filed by Oxford University Innovations for single-cell technologies.

