## Supplemental Figures for "Highly accurate barcode and UMI error correction using dual nucleotide dimer blocks allows direct single-cell nanopore transcriptome sequencing"

Supplementary Figures

Figure 1

­­
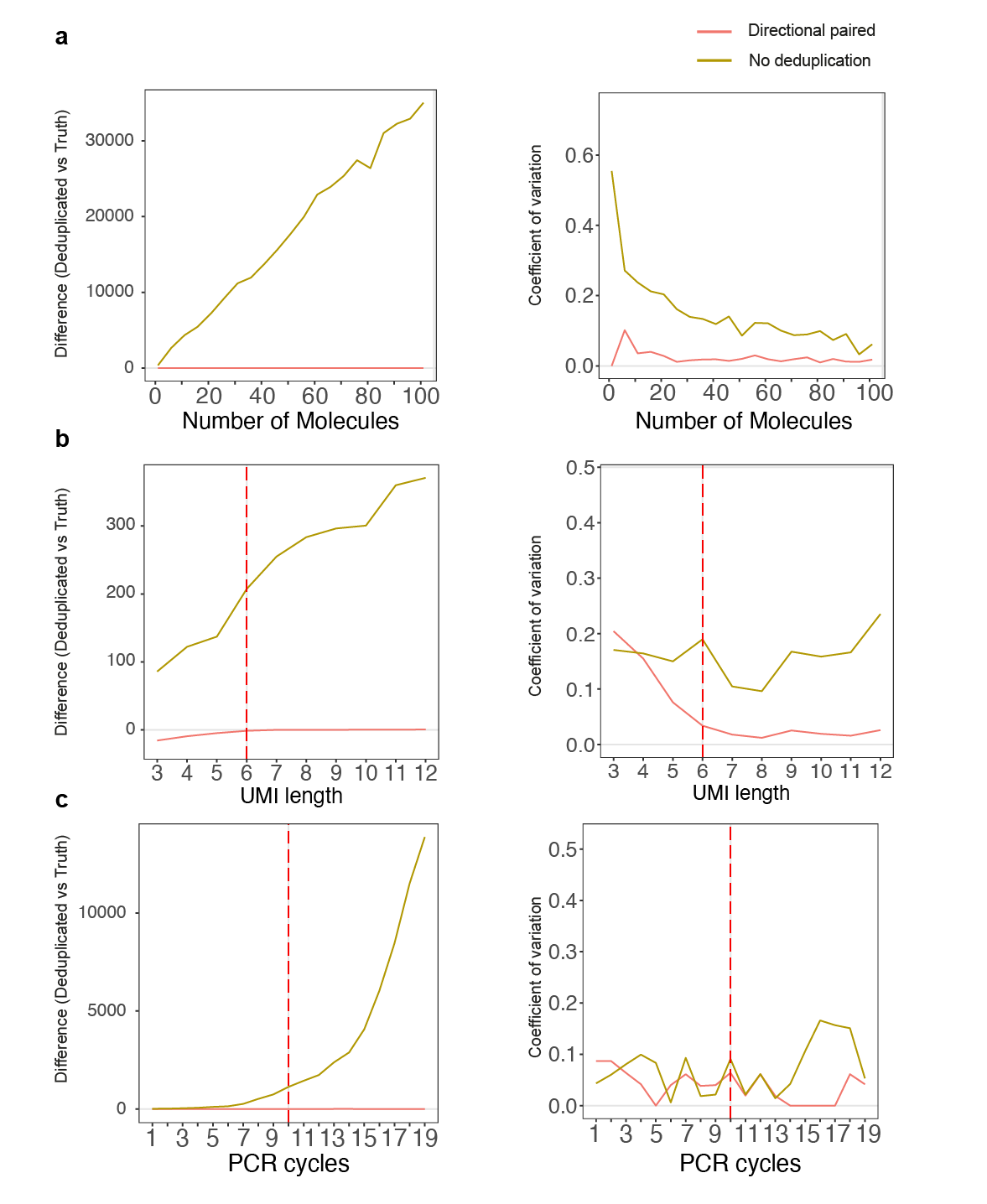

**Simulated data showing the application of directional UMI deduplication strategy**

Simulated data showing the difference and coefficient of variation between deduplicated UMIs and the ground truth. The left-hand pane shows the difference between the method deduplication UMI numbers and the simulated ground truth. The right-hand pane shows the coefficient of variation following 10 iterations. **a** The effect of increasing the starting number of molecules on the ability to deduplicate UMIs. **b** The effect of increasing the UMI length on the ability to deduplicate UMIs. **c** The effect of increasing the number of simulated PCR cycles using an error rate of 1x10^-5^.

Figure 2

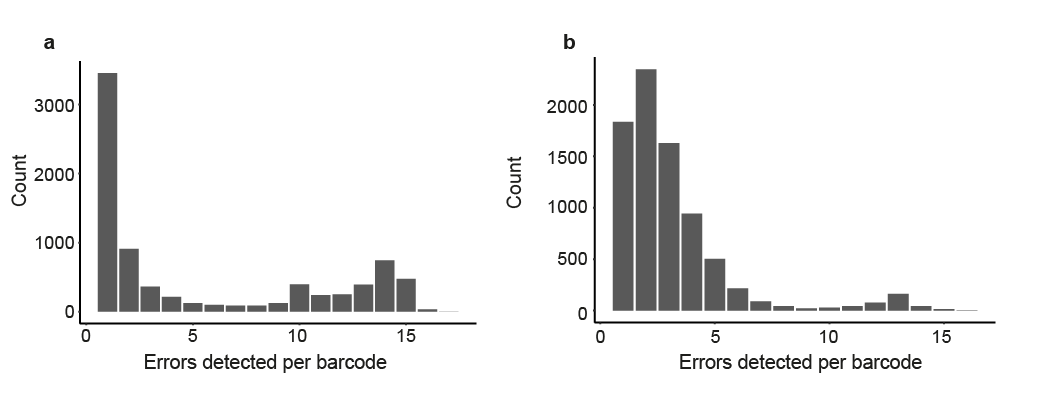

**The frequency of errors within barcodes that contain at least one error.**

HEK and 3T3 cells were encapsulated and then a library was prepared for both Illumina and Nanopore sequencing using the same cDNA**. a** 8000 Illumina sequenced barcodes were randomly selected from barcodes that contained at least one sequencing error. The frequency of sequencing errors is plotted as a bar graph. **b** Similar for **a**, but for nanopore sequenced barcodes.

Figure 3

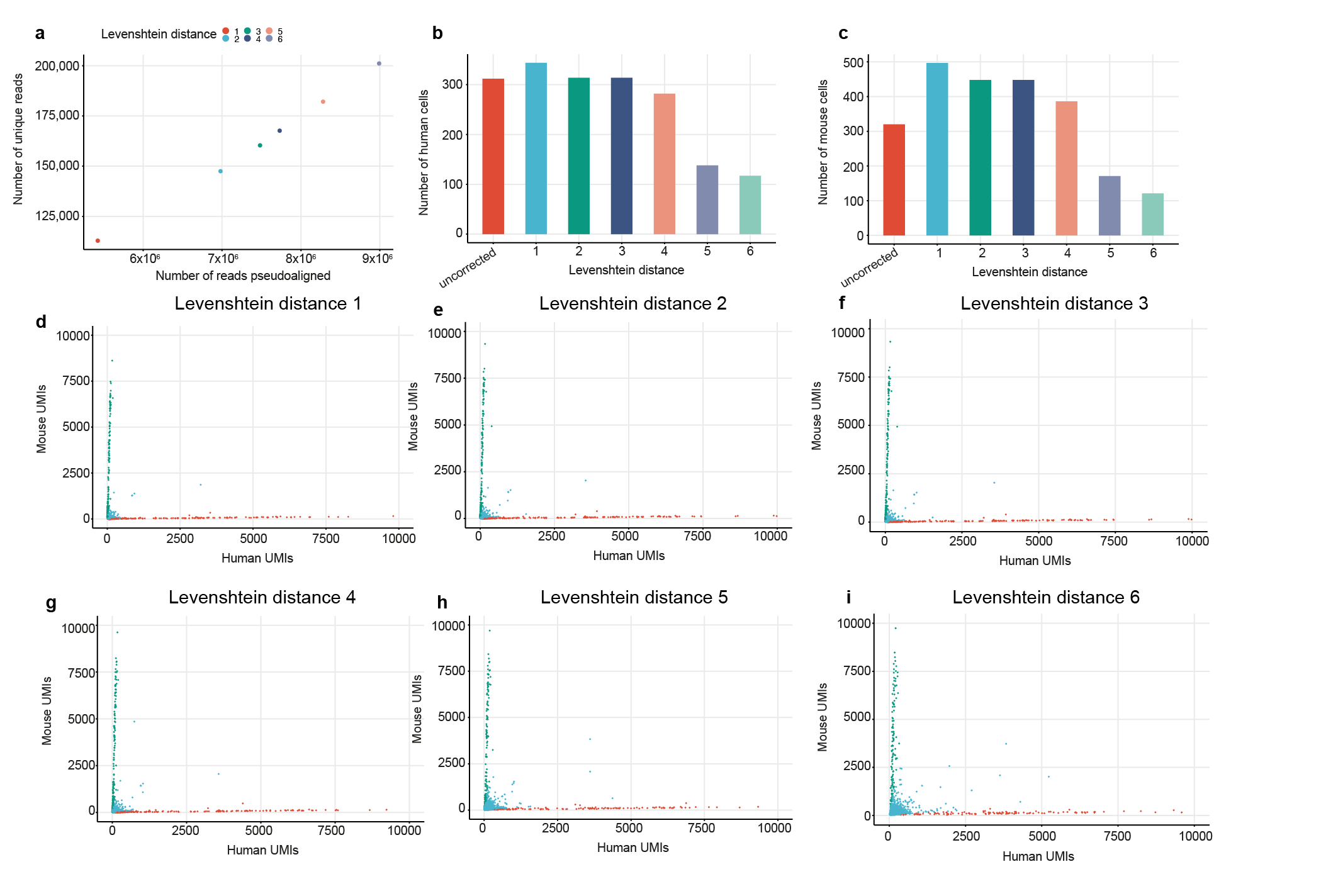

**Evaluating different edit distances for correcting Illumina scRNA sequencing data.**

A dual oligonucleotide scRNA-seq library was generated and around 500 human HEK293T and mouse 3T3 cells were sequenced using the Illumina platform. Barcodes that contained a sequencing error, as determined by dual nucleotide block complementarity were identified. Barcodes were then error corrected using increasing edit distances. **a** The relationship between the number of unique pseudoaligned reads compared to the number of pseudoaligned reads following barcode error correction with increasing Levenshtein distance. **b** The number of human cells identified using increasing Levenshtein distance for barcode error correction. **c** The corresponding numbers of mouse cells identified with increasing Levenshtein distance. **d, e, f, g, h, i** Barnyard plots showing mouse and human UMIs detected per cell.

Figure 4

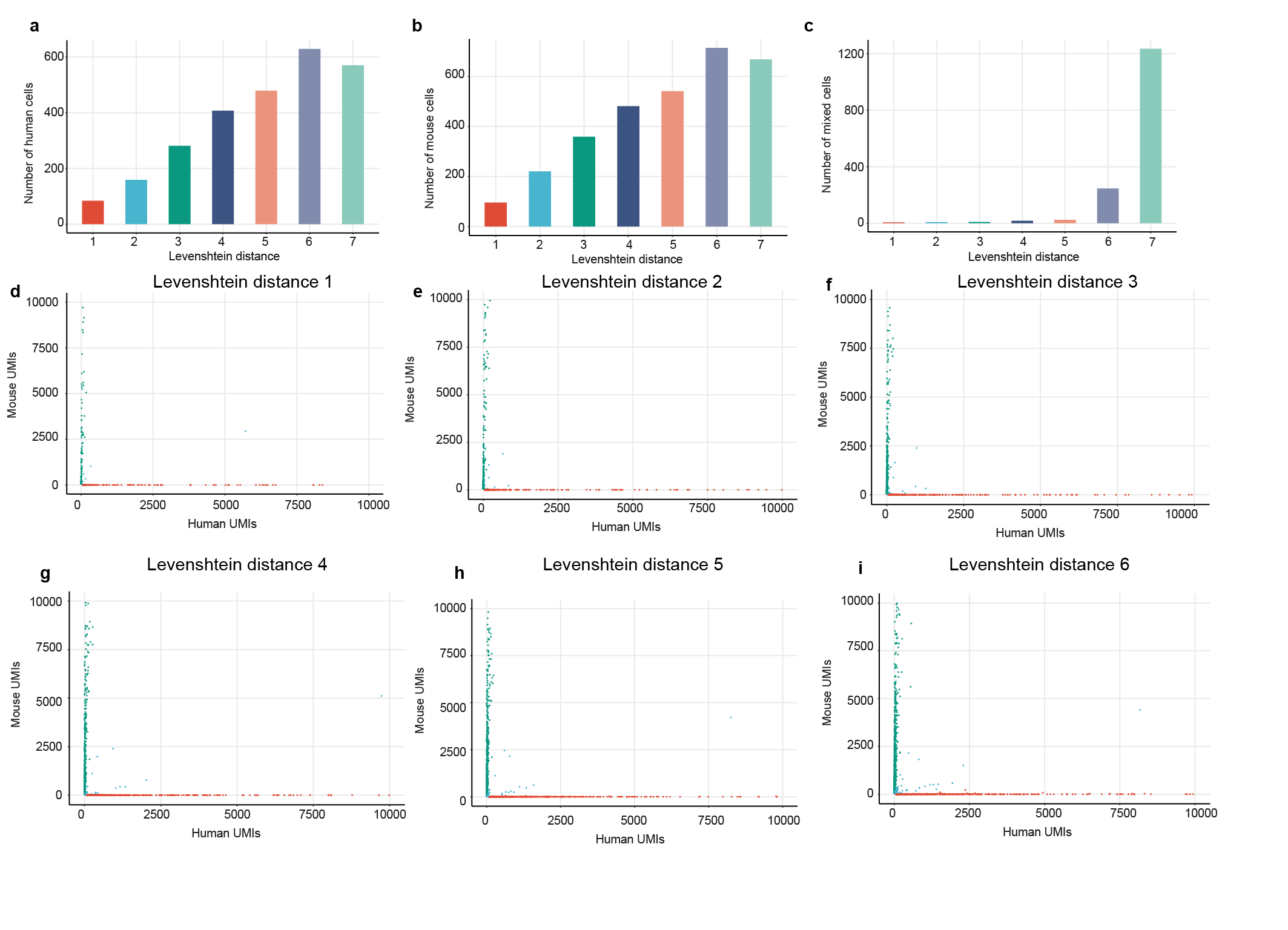

**Evaluating different edit distances for correcting Nanopore scRNA sequencing data.**

A dual oligonucleotide scRNA-seq library was generated and around 500 human HEK293T and mouse 3T3 cells were sequenced using the Oxford Nanopore platform. Barcodes that contained a sequencing error, as determined by dual nucleotide block complementarity were identified. Barcodes were then error corrected using increasing edit distances. **a** The number of human cells identified using increasing Levenshtein distance for barcode error correction. **b** The corresponding numbers of mouse cells identified with increasing Levenshtein distance. **c** The corresponding numbers of mixed cells identified with increasing Levenshtein distance.

**d, e, f, g, h, i** Barnyard plots showing mouse and human UMIs detected per cell.

Figure 5

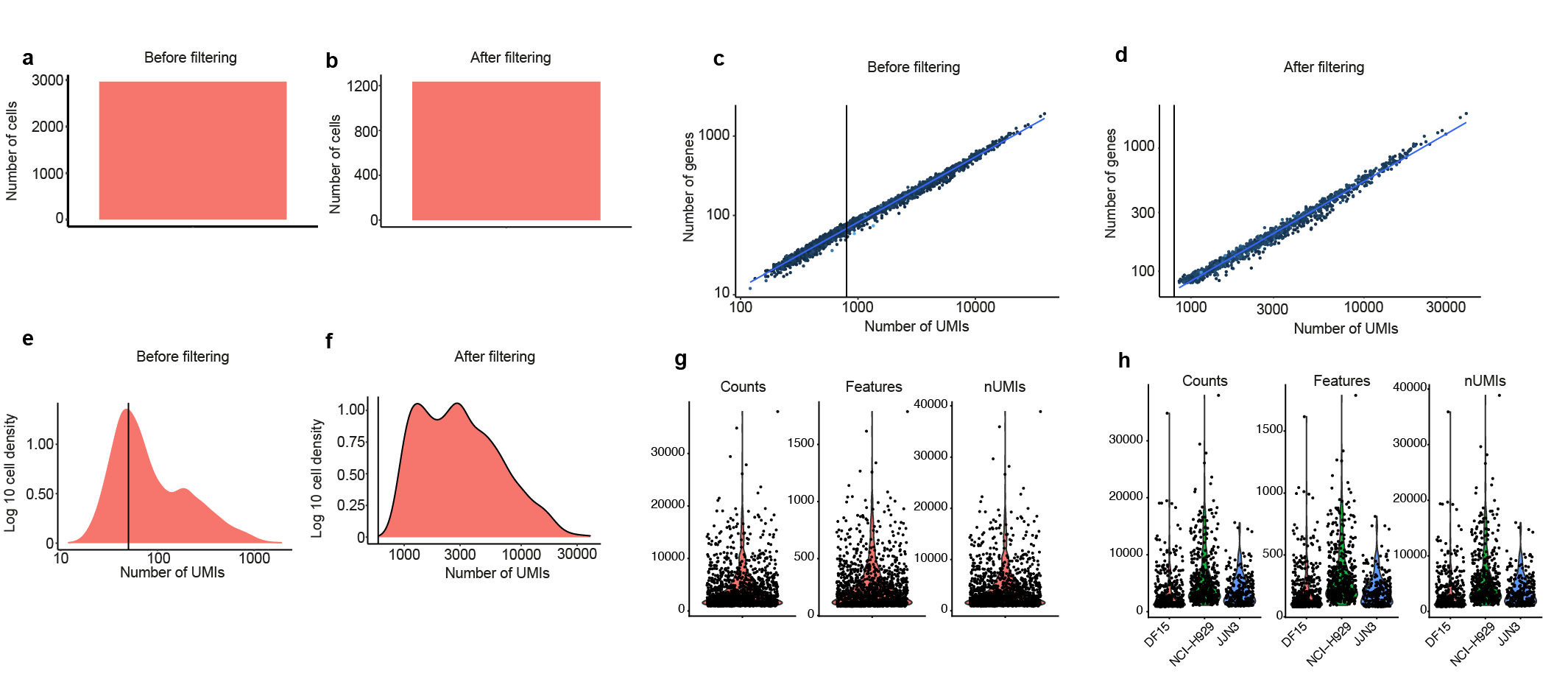

**Filtering and removal of low-quality cells from 1200 NCI-H929, JJN3 and DF15 mixed cell experiment sequenced using a PromethION.**

Cells expressing greater than 600 UMIs and 80 genes per cell was used as a threshold to filter poor quality cells from our 1200-cell mixed myeloma cell line dataset. The number of cells **a** before and **b** after filtering. The relationship between the number of UMIs and the number of genes **c** before and **d** after filtering. ­A histogram of the number of UMIs **e** before and **f** after filtering. The number of counts, features and UMIs across **g** all filtered cells and across each **h** myeloma cell type.

Figure 6

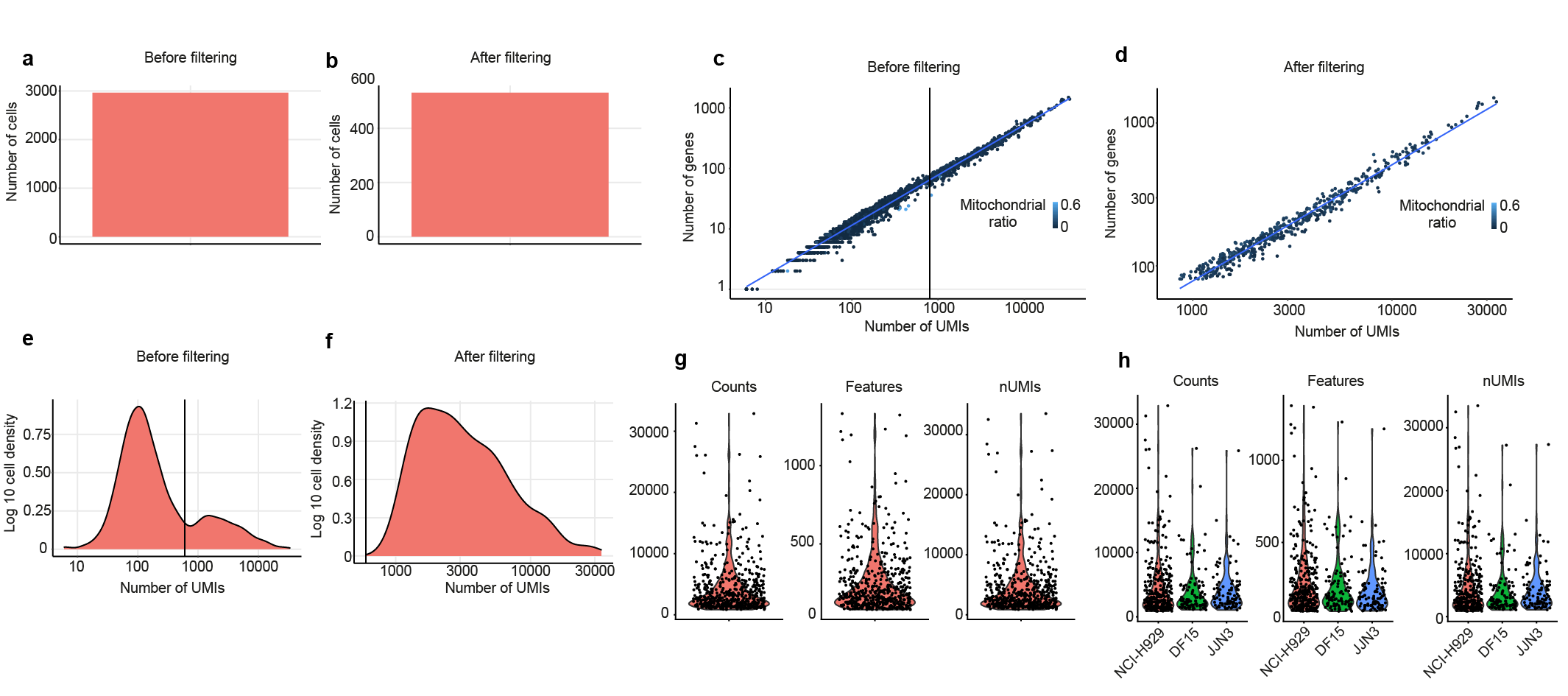

**Filtering and removal of low-quality cells from 500 NCI-H929, JJN3 and DF15 mixed cell experiment sequenced using a MinION.**

Cells expressing greater than 600 UMIs and 80 genes per cell was used as a threshold to filter poor quality cells from our 500-cell mixed myeloma cell line dataset. The number of cells **a** before and **b** after filtering. The relationship between the number of UMIs and the number of genes **c** before and **d** after filtering. ­A histogram of the number of UMIs **e** before and **f** after filtering. The number of counts, features and UMIs across **g** all filtered cells and across each **h** myeloma cell type.

Figure 7

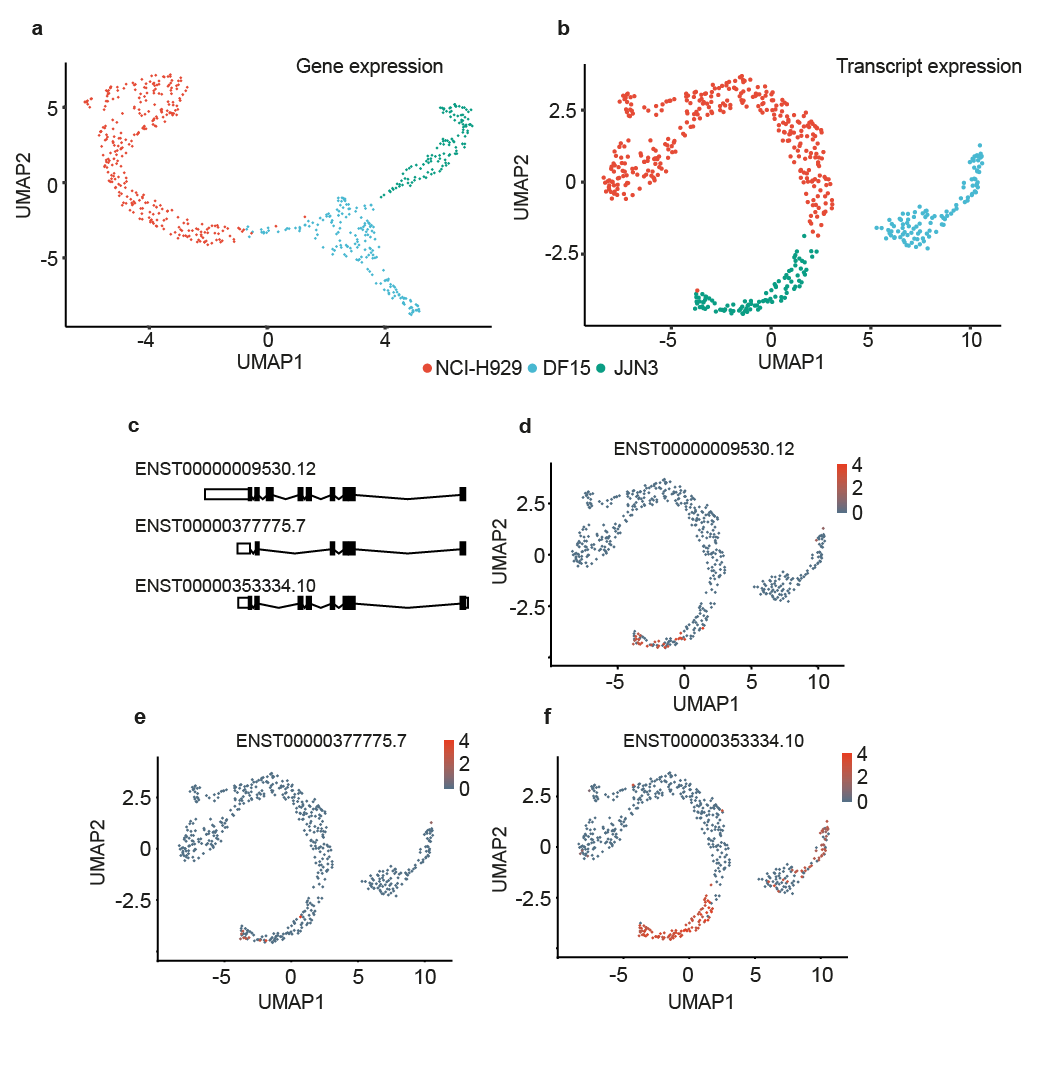

**Nanopore droplet based scRNA-seq identifies isoform diversity in the 500 myeloma MinION sequenced experiment.**

NCI-H929, DF15 and JJN3 myeloma cell lines were mixed at a 1:1:1 ratio and approximately 500 cells were taken for cDNA synthesis and sequenced using a MinION flow cell. UMAP plot of **a** gene expression and **b** transcript isoform expression. **c** Principal CD74 (HLA-DR) splice variants showing all protein coding transcripts. UMAP plot showing the isoform expression of detected CD74 (HLA-DR) transcripts **d** ENST00000009530.12, **e** ENST00000377775.7 and **f** ENST00000353334.10.

Figure 8

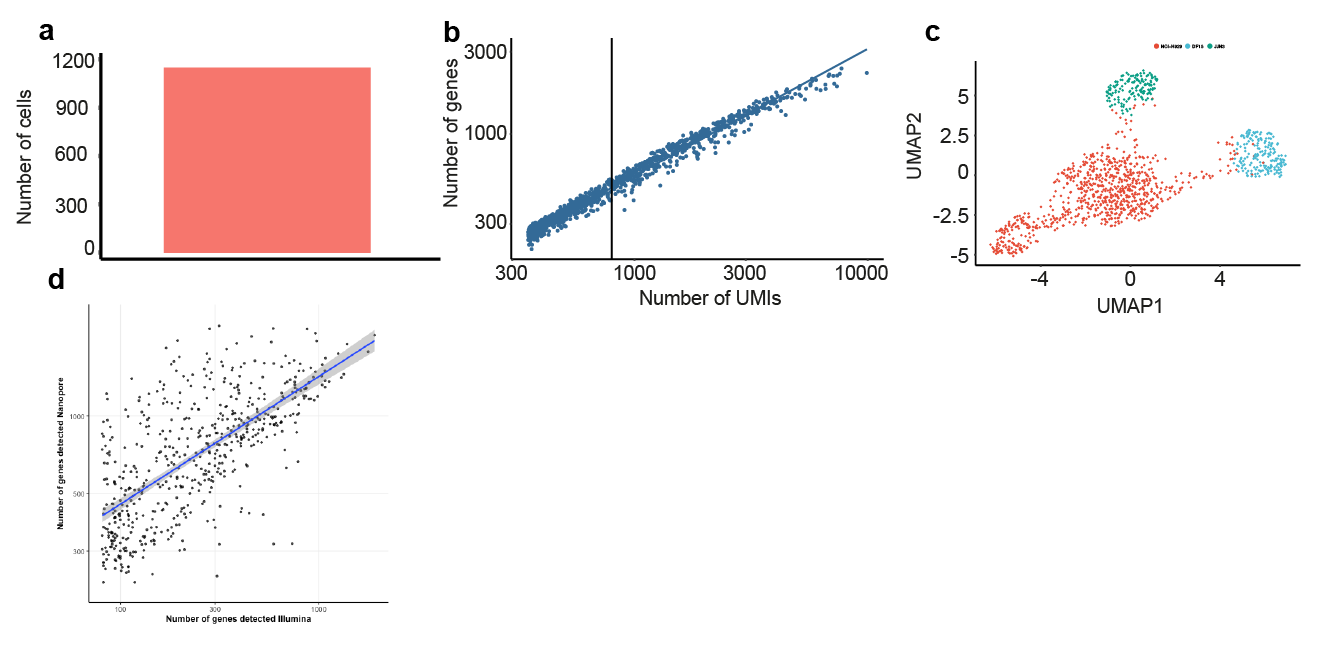

**Correlation between Illumina and Nanopore sequencing**

NCI-H929, DF15 and JJN3 myeloma cell lines were mixed at a 1:1:1 ratio and approximately 1200 cells were taken for cDNA synthesis (The same library that was sequenced in Figure 3) and sequenced using the Illumina platform. Cells expressing greater than 300 genes per cell was used as a threshold to filter poor quality cells. **a** The number of cells after filtering. **b** The relationship between the number of UMIs and the number of genes after filtering. **c** UMAP plot of gene expression. **d** The correlation between the number of genes detected by Illumina sequencing and Nanopore sequencing.
Figure 9

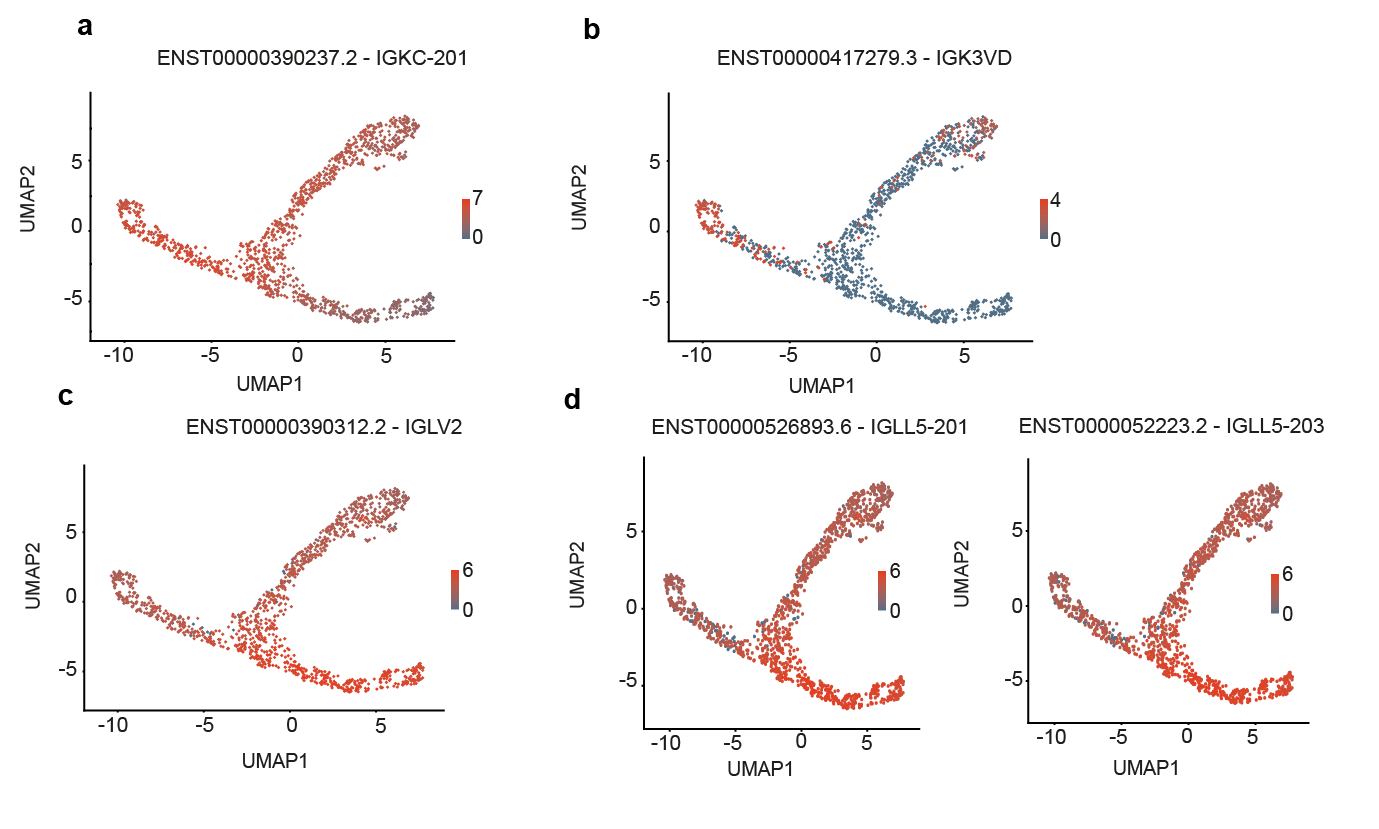

**The expression of Immunoglobulin Kappa and Lambda constant transcripts in the 1200 cell myeloma experiment.**

UMAP plot showing the expression of **a** IGKC-201, **b** IGKV3D-15, **c** IGLV2 and **d** IGLL5.

Figure 10

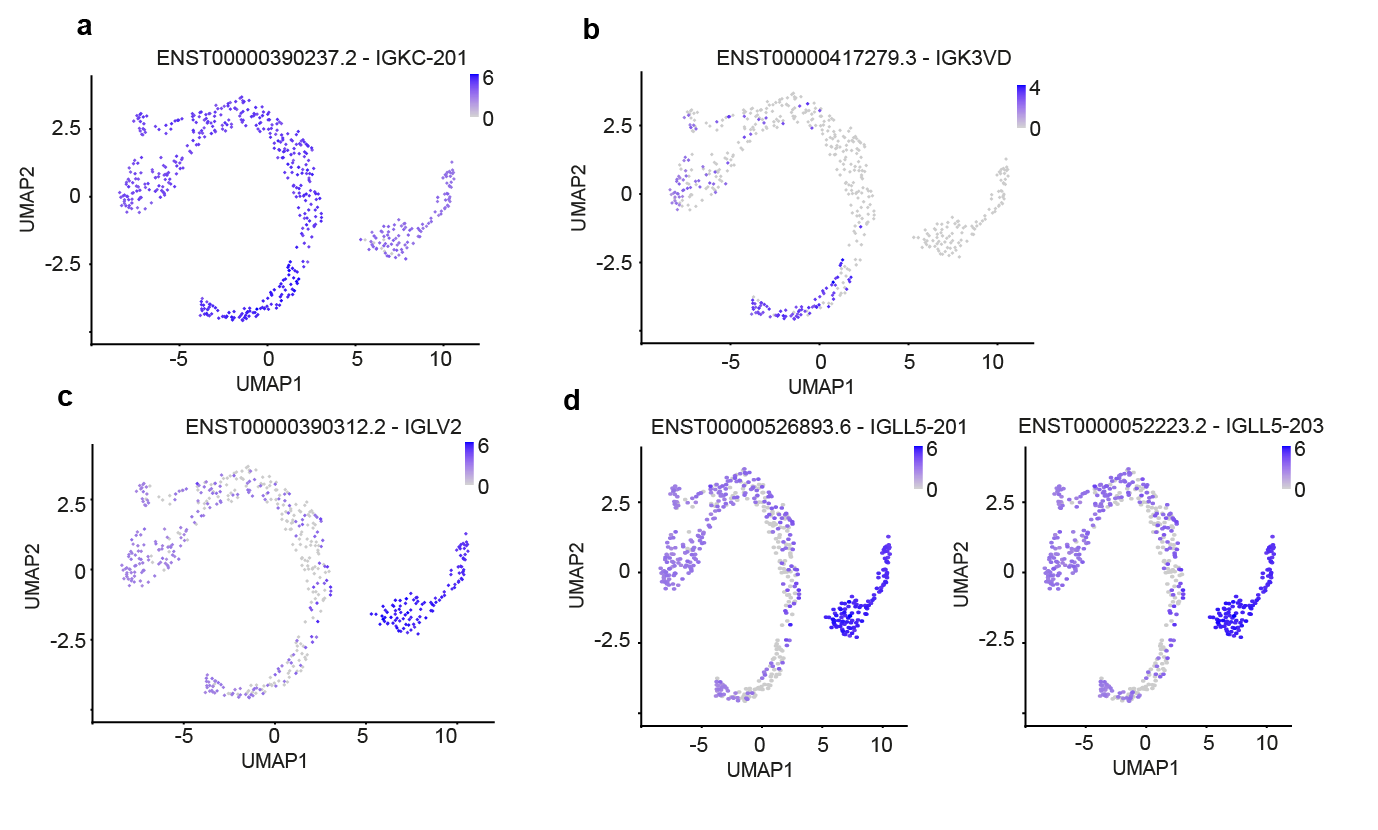

**The expression of Immunoglobulin Kappa and Lambda constant transcripts in the 500 cell myeloma experiment.**

UMAP plot showing the expression of **a** IGKC-201, **b** IGKV3D-15, **c** IGLV2 and **d** IGLL5.

Figure 11

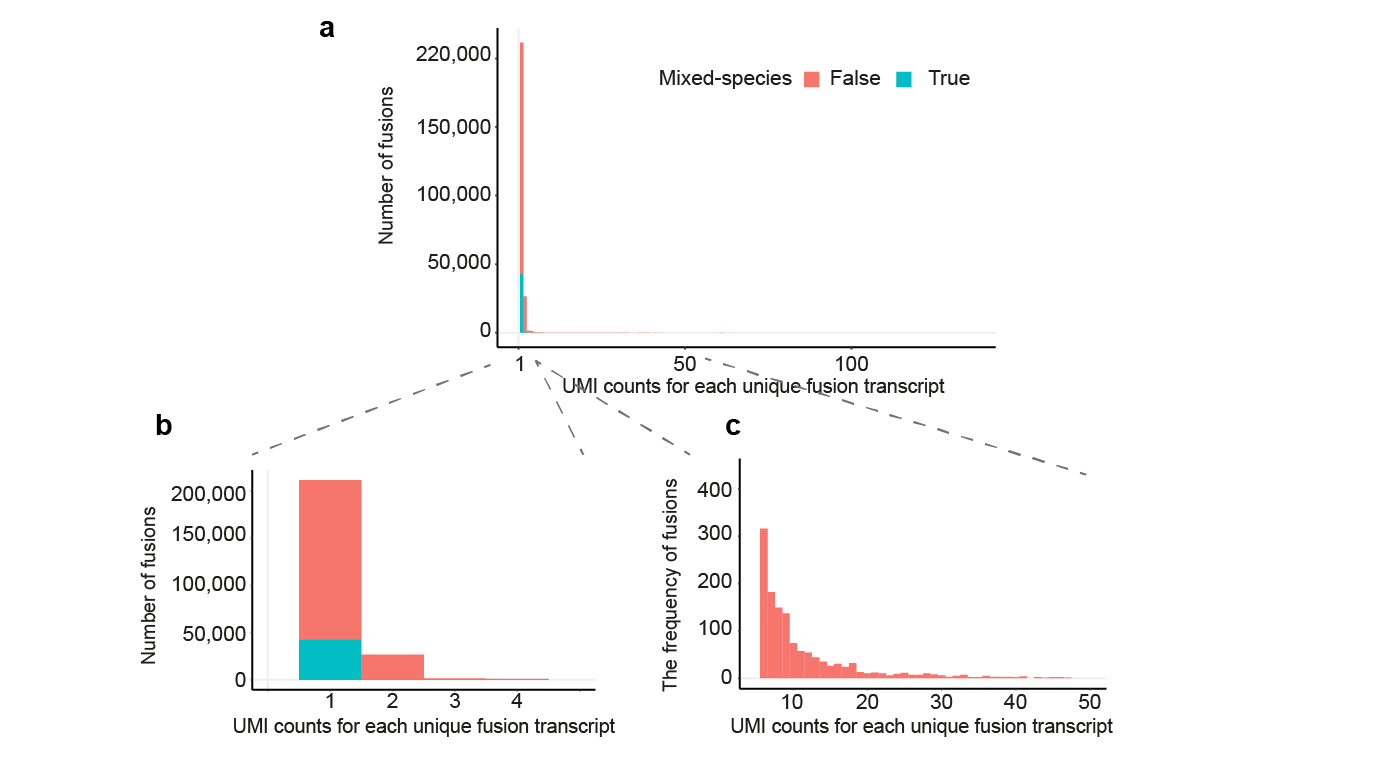

**The detection of fusion transcripts within the mixed species dataset**

We measured the presence of fusion transcripts within our mouse and human mixed species experiment. **a** The frequency of UMI counts per unique fusion transcript. b The same data shown in **a,** but limited to UMI counts between 1 and 5. **c** The same data shown in **a** but limited to UMI counts between 5 and 50. The colours indicate the presence of mixed species.
Figure 12

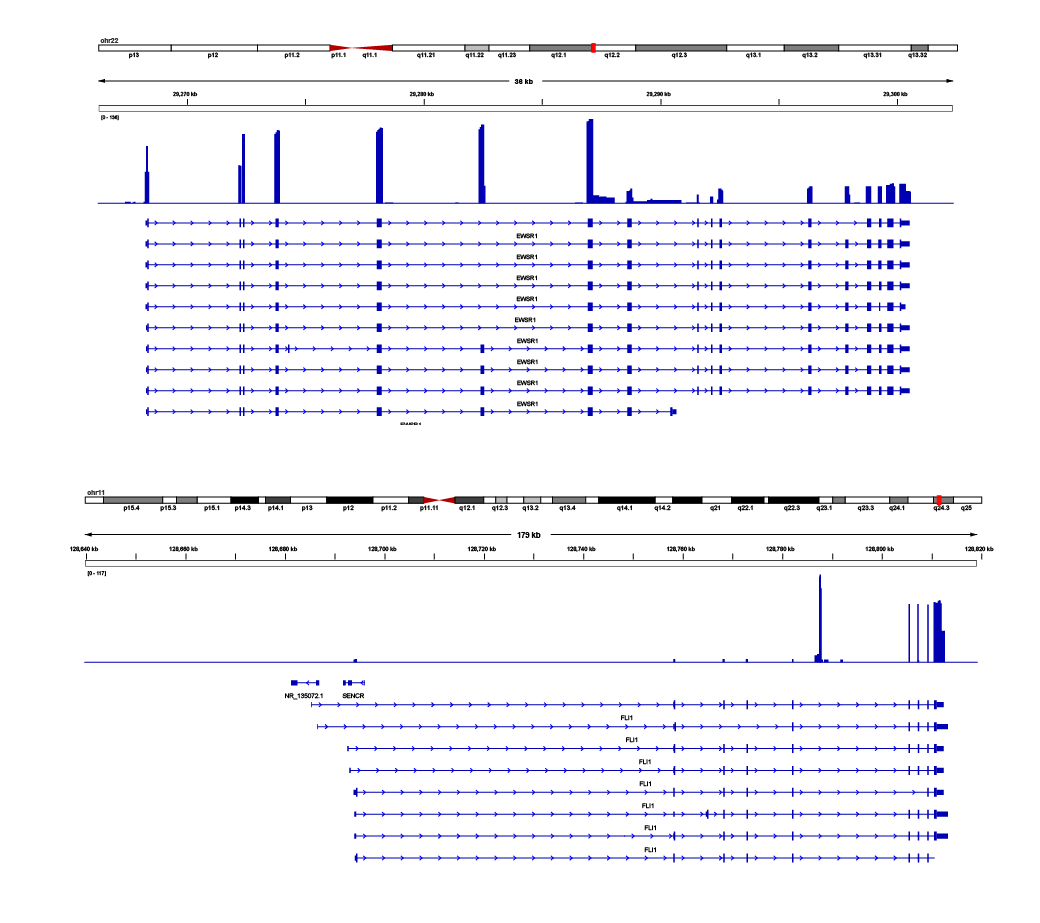

**Genome browser tracks showing the read pile up across both EWSR1 and FLI1**

The top panel shows the read pileup across the exons of EWSR1. The bottom panel shows the read pileup across the FLI1 gene. The peak observed within the intronic region between exons 5 and 6 appears to be an alignment artifact and does not represent a real peak.
